# *Cbs* overdosage is necessary and sufficient to induce cognitive phenotypes in mouse models of Down syndrome and interacts genetically with *Dyrk1a*

**DOI:** 10.1101/393579

**Authors:** Damien Marechal, Véronique Brault, Alice Leon, Dehren Martin, Patricia Lopes Pereira, Nadege Loaёc, Marie-Christine Birling, Gaelle Friocourt, Marc Blondel, Yann Herault

**Affiliations:** Institut de Génétique et de Biologie Moléculaire et Cellulaire, Illkirch, 1 rue Laurent Fries, 67404 Illkirch, France; Centre National de la Recherche Scientifique, UMR7104, Illkirch, France; Institut National de la Santé et de la Recherche Médicale, U1258, Illkirch, France; Université de Strasbourg, Illkirch, France; Inserm UMR 1078, Université de Bretagne Occidentale, Faculté de Médecine et des Sciences de la Santé, Etablissement Français du Sang (EFS) Bretagne, CHRU Brest, Hôpital Morvan, Laboratoire de Génétique Moléculaire, Brest, France; Transgenese et Archivage Animaux Modèles TAAM, CNRS, UPS44, 3B rue de la Férollerie 45071 Orléans, France; CELPHEDIA, PHENOMIN, Institut Clinique de la Souris, ICS, 1 rue Laurent Fries, 67404 Illkirch, France

## Abstract

Identifying dosage sensitive genes is a key to understand the mechanisms underlying intellectual disability in Down syndrome (DS). The Dp(17Abcg1-Cbs)1Yah DS mouse model (Dp1Yah) show cognitive phenotype and needs to be investigated to identify the main genetic driver. Here, we report that, in the Dp1Yah mice, 3 copies of the Cystathionine-beta-synthase gene (*Cbs)* are necessary to observe a deficit in the novel object recognition (NOR) paradigm. Moreover, the overexpression of *Cbs* alone is sufficient to induce NOR deficit. Accordingly targeting the overexpression of human CBS, specifically in Camk2a-expressing neurons, leads to impaired objects discrimination. Altogether this shows that *Cbs* overdosage is involved in DS learning and memory phenotypes. In order to go further, we identified compounds that interfere with the phenotypical consequence of CBS overdosage in yeast. Pharmacological intervention in the Tg(*CBS*) with one selected compound restored memory in the novel object recognition. In addition, using a genetic approach, we demonstrated an epistatic interaction between *Cbs* and *Dyrk1a*, another human chromosome 21 gene encoding the dual-specificity tyrosine phosphorylation-regulated kinase 1a and an already known target for DS therapeutic intervention. Further analysis using proteomic approaches highlighted several pathways, including synaptic transmission, cell projection morphogenesis, and actin cytoskeleton, that are affected by DYRK1A and CBS overexpression. Overall we demonstrated that CBS overdosage underpins the DS-related recognition memory deficit and that both *CBS* and *DYRK1A* interact to control accurate memory processes in DS. In addition, our study establishes CBS as an intervention point for treating intellectual deficiencies linked to DS.

**SIGNIFICANT STATEMENT:** Here, we investigated a region homologous to Hsa21 and located on mouse chromosome 17. We demonstrated using three independent genetic approaches that the overdosage of the Cystathionine-beta-synthase gene (*Cbs*) gene, encoded in the segment, is necessary and sufficient to induce deficit in novel object recognition (NR).

In addition, we identified compounds that interfere with the phenotypical consequence of CBS overdosage in yeast and in mouse transgenic lines. Then we analyzed the relation between Cbs overdosage and the consequence of DYRK1a overexpression, a main driver of another region homologous to Hsa21 and we demonstrated that an epistatic interaction exist between *Cbs* and *Dyrk1a* affecting different pathways, including synaptic transmission, cell projection morphogenesis, and actin cytoskeleton.

## INTRODUCTION

Down Syndrome (DS) is the most common aneuploidy observed in human. The presence of an extra copy of the Human chromosome 21 (Hsa21; Hsa for *Homo sapiens*) is associated with intellectual disabilities and several morphological and physiological features. Phenotypic mapping in human with partial duplication highlighted the contribution of several regions of the Hsa21 in DS features (1, 2). Additional information was collected from trisomic and monosomic mouse models to detect genomic regions sensitive to dosage and able to induce impairments in behaviour and other DS related traits (3-11). Most of the efforts focused on the region homologous to the Hsa21 located on mouse chromosome 16 (Mmu16; *Mmu for Mus Musculus*), highlighting the contribution of the Amyloid precursor protein (*App*) (12), of the Glutamate receptor, ionotropic, kainate 1 (*Grik1)* or of the dual-specificity tyrosine phosphorylation-regulated kinase 1a *(Dyrk1a)* (13, 14) overdosage to DS cognitive defects. At present, DYRK1A is a main target for therapeutic intervention with a few compounds inhibiting the protein kinase activity, improving mainly cognition in DS mouse models (15-20). However, models carrying trisomy of the region of Mmu17 homologous with the Hsa21, also showed learning and memory defects (21, 22) and appeared to have a major impact on DS phenotypes in mouse models (23). The Dp(17*Abcg1-Cbs*)1Yah (called here Dp1Yah) mice are defective in the novel object recognition test and show a long-lasting in vivo long-term potentiation (LTP) in the hippocampus while the corresponding monosomy, Ms2Yah, have defects in social discrimination with increased in vivo LTP (24). Interestingly, as observed in the rotarod test, the locomotor phenotype of the Tc1 transchromosomic model carrying an almost complete Hsa21 is rescued when the dosage of the *Abcg1-Cbs* region is reduced in Tc1/Ms2Yah mice (25). Similarly the trisomy of a larger overlapping segment on Mmu17 from *Abcg1* to *Rrp1b* induces an increased LTP as compared to control in the Dp(17)Yey model (22) and was shown to genetically interact with the trisomy of the *Lipi-Zbtb21* interval. More specifically the trisomy of both this *Abcg1*-*Rrp1b* region and the *Cbr1-Fam3b* region was detrimental for learning and memory in the Morris water maze and for LTP in DS mouse models (23).

Among the 11 trisomic genes in the Dp1Yah model, the cystathionine-beta-synthase gene, *Cbs*, encodes a pyridoxal phosphate-dependent enzyme converting homocysteine to cystathionine. This first step of the transulfuration pathway removes homocysteine from the methionine cycle thereby also affecting the folate and the methylation pathways, while contributing to the cysteine cycle. Of note, in human, homozygous loss-of-function mutations in *CBS* are associated with homocystinuria (OMIN236200) a metabolic condition with intellectual disability. CBS is also the major enzyme catalysing the production of H2S from L-cysteine (26) or from the condensation of homocysteine with cysteine (27). H2S is now considered a major gaseotransmitter in the brain (28) and interferes with synaptic transmission. Considering the upregulated expression of CBS in several brain regions of the Dp1Yah model and its impact on intellectual disability, we decided to focus on *Cbs* and decipher the role of CBS in DS cognitive phenotypes. To this end, we generated and characterized constitutive and conditional changes in *Cbs* dosage in the nervous system of various mouse models. In addition we selected pharmacological drugs able to counteract the phenotypical consequence of CBS overexpression, in particular behavioural impairments, and finally further analysed molecular changes induced by *Cbs* dosage changes to understand the mechanisms perturbed in DS models.

## MATERIALS AND METHODS

### Ethics Statement, mouse lines and genotyping

Animal experiments were approved by the Com’Eth N°17 (project file: 2012-069) and accredited by the French Ministry for Superior Education and Research and in accordance with the Directive of the European Parliament: 2010/63/EU, revising/replacing Directive 86/609/EEC and with French Law (Decret n° 2013-118 01 and its supporting annexes entered into legislation 01 February 2013) relative to the protection of animals used in scientific experimentation. YH was granted the accreditation 67-369 to perform the reported experiments in the animal facility (Agreement C67-218-40). For all these tests, mice were kept in Specific Pathogen free conditions with free access to food and water. The light cycle was controlled as 12 h light and 12 h dark (lights on at 7AM). All the behavioural tests were done between 9:00 AM and 4:00 PM.

Several mouse lines were used to decipher the influence of *Cbs*: the trisomic mouse model, Dp(17Abcg1-Cbs)1Yah, named here Dp1Yah, carries a segmental duplication of the *Abcg1-Cbs* region of the Mmu17 (21) kept on the C57BL/6J; the inactivated allele of C57BL/6J*.Cbs^tm1Unc^* (29); and the PAC transgenic line Tg(*CBS*)11181Eri (named here Tg(*CBS*)), originally identified as 60.4P102D1 (30) and backcrossed on C57BL/6J for more than 7 generations. We designed, generated and selected the transgenic mouse line Tg(*Prp-gfp-CBS*)95-157ICS, named here Tg(*Prp-gfp-CBS*), to overexpress the human *CBS* cDNA from the murine prion promoter region (containing a 8477 bp region upstream of the ATG of the murine prion gene, ie 6170 bp promoter region, exon1, intron 1 and beginning of exon 2) after the excision of a loxP-*gfp-loxP* interrupting cassette (Figure 3A) on C57BL/6J background. We used the transgenic Tg(Camk2a-cre)4Gsc mouse line (31), named here Tg(*Camk2a-cre*), and bred further on C57BL/6J, as a glutamatergic neuron-specific Cre driver. The Dyrk1a BAC transgenic mouse line, named here Tg(*Dyrk1a*) was generated previously in our lab (32). All lines were generated and bred on the C57BL/6J genetic. The genotype identification was done from genomic DNA isolated from tail biopsies with specific PCR reaction (Supplementary table 1).

**Figure 3.**
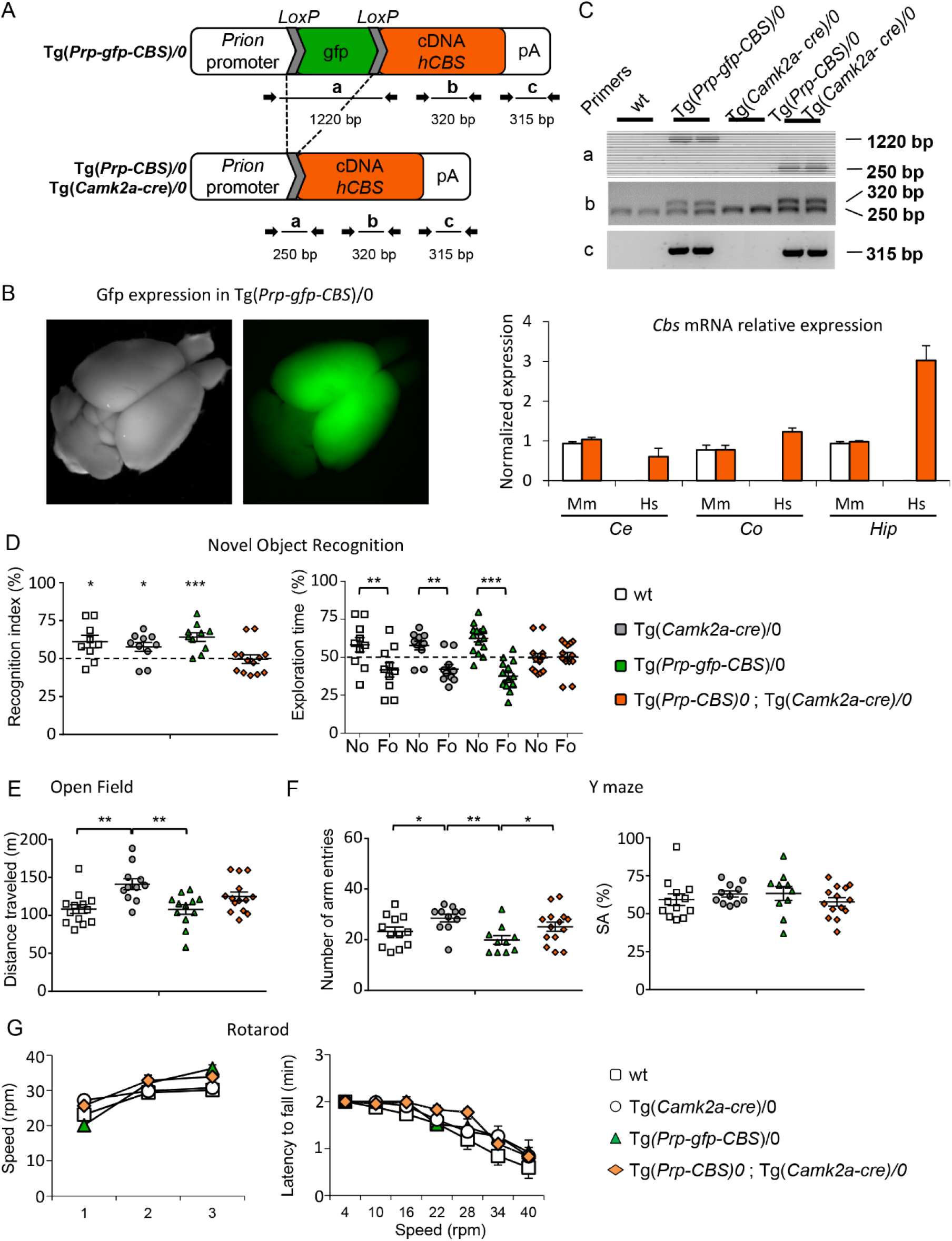
Selective overexpression of *hCBS* in the glutamatergic neurons leads to impaired object recognition and altered locomotor activity. (A) a conditional transgene Tg(*Prp-gfp-CBS*) was designed to overexpress the human CBS cDNA from the murine Prion promoter after the deletion of an interrupting GFP-coding cassette flanked by loxP sites. The GFP allowed to select one line that lead to expression in the anterior part of the brain (B). When Cre is expressed from the Tg(Camk2-Cre) transgene, the deletion can be monitored in the brain of the animals (C) and the overexpression of hCBS mRNA is detected in different part of the brain of the Tg(Camk2-Cre)/0;Tg(*Prp-gfp-CBS*)/0 animals (Hs, orange bar, B) with no change in the endogeneous murine CBS without Cre expression detected in wt animals (Mm, white bar B) or Tg(Camk2-Cre)/0;Tg(*Prp-gfp-CBS*)/0 animals (Mm, orange bar, B). Wt (n=13), Tg(Camk2-Cre)/0 (n=11), Tg(*Prp-gfp-CBS*)/0 (n=12), and Tg(Camk2-Cre)/0;Tg(*Prp-gfp-CBS*)/0 (n=14) littermates were evaluated through for object discrimination (D), open field (E), Y maze (F), rotarod (G). Mice overexpressing hCBS in the glutamatergic neurons were unable to discriminate the novel versus the familiar object as compared to the other control genotypes (D). Tg(Camk2-Cre)/0 mice displayed an enhanced locomotor activity in the open field but no change was detected in the control, wt and Tg(*Prp-gfp-CBS*)/0, or in double transgenic animals (E). In the Y maze animals carrying the Tg(*Prp-gfp-CBS*)/0 or the activated form, Tg(Camk2-Cre)/0;Tg(*Prp-gfp-CBS*)/0, displayed reduced exploration with a lower number of arm entries but no change in the spontaneous alternation (F). No phenotypes was altered in the rotarod test with similar progress during the learning and the test phases (G). (Values represent means + S.E.M. *P<0.05, **P<0.01, ***P<0.001)

### Behavioural analysis

The sample size was estimated according to our similar experiments done previously while investigating behaviour in DS mouse models (5, 25, 33). To investigate the role of *Cbs* in the Dp1Yah cognitive phenotypes, we generated 2 independent cohorts (cohort 1 (C1): wild type (wt) littermates n=11; *Cbs^tm1Unc/+^*, n=8; Dp1Yah, n=8; Dp1Yah/*Cbs^tm1Unc^*, n=11; and cohort 2 (C2): wt littermates n=18; *Cbs^tm1Unc/+^*, n*=*15; Dp1Yah, n=15; Dp1Yah/*Cbs^tm1Unc^*, n=10). All cohorts were evaluated in the open field (C1: 33 weeks; C2: 14-16 weeks), Novel Object Recognition (NOR) (C1: 33 weeks; C2:14-16 weeks) in adult mice. In addition we performed the Y maze (C2: 15-19 weeks) and the rotarod tests (C2: 25-28 weeks of age).

Wild-type littermates (n=13) and Tg(CBS)/0 (n=17) hemizygotes were tested for circadian actimetry (14 weeks), Y Maze (16 weeks), open field (17 weeks) and NOR (17 weeks). We added an additional group of wt (n=9) and Tg(CBS)/0 (n=10) to validate the results from the NOR; animals were tested at the same age (17 weeks). A cohort with 4 genotypes (wt (n=13), Tg(Camk2-Cre)/0 (n=11), Tg(*Prp-gfp-CBS*)/0 (n=12), and Tg(Camk2-Cre)/0;Tg(*Prp-gfp-CBS*)/0 (n=14)) was evaluated through the same behavioural tests with rotarod (14 weeks), Y maze (16 weeks), open field (19-20 weeks) and NOR (19-20 weeks). 14 wt, 15 Tg(*Dyrk1a*), 13 Dp1Yah and 13 Dp1Yah/Tg(*Dyrk1a*) mutant mice were evaluated for open field exploration (11-12 weeks), novel object recognition (11-12 weeks) and Y maze (13 weeks). A second independent cohort with 11 wt, 10 Tg(*Dyrk1a*), 14 Dp1Yah and 10 Dp1Yah/Tg(*Dyrk1a*) was used for Morris water maze learning (14-16 weeks). The behavioural protocols for open-field, Y maze and novel object recognition, rotarod, water maze were are detailed in the supplementary information.

### Drug screening in yeast

All plasmids were generated using standard procedures. Restriction enzymes and Taq polymerase were obtained from New England Biolabs (Evry, France). T4 DNA ligase was purchased from Promega and purified synthetic oligonucleotides from Eurogentec. Routine plasmid maintenance was carried out in DH5α and TOP10 bacteria strains. Yeast cystathionine b-synthase (Cys4) coding sequence was amplified from the genomic DNA of the W303 *WT* strain (see genotype below) using Bam-Cys4-F: CGGGATCCCGATGACTAAATCTGAGCAGCAAG and Xho-Cys4-R: GCCTCGAGTCTTATGCTAAGTAGCTCAGTAAATCC (that introduced *Bam*HI and *Xho1* restriction sites) and subcloned in the high copy number 2 μ-derived vectors p424-GPD and p426-GPD, each time under the control of the strong constitutive *GDP* promoter (34). Transformation of yeast cells was performed using a standard lithium acetate method (35).

The yeast strain used in this study is derived from the W303 *WT* strain: *MATa*, *leu2-3,112 trp1-1 can1-100 ura3-1 ade2-1 his3-11,15*. The media used for yeast growth were: YPD [1% (w/v) yeast extract, 2% (w/v) peptone, 2% (w/v) glucose, for untransformed cells and Synthetic Dextrose *Minimal medium* (SD medium) (composed of 0.67% (w/v) *Yeast* Nitrogen Base w/o amino acids and complemented with 0.1% (w/v) casamino acid, 40 mg/l adenine and 2% (v/v) glucose for Cys4-transformed cells. Solid media contained 2% (w/v) agar.

For the drug screening, yeast cells were grown in uracil- and tryptophan-free minimal liquid medium (SD-Ura/Trp) in overnight liquid cultures at 29 °C. The following day, cells were diluted to OD600~0.2 in in fresh medium and grown for 4 hours to reach exponential phase. Then three hundred and fifty microliters of exponentially growing yeast cells overexpressing Cys4, adjusted to an OD600 of 0.5, were spread homogeneously with sterile glass beads (a mix of ~1.5 and 3 mm diameter) on a square Petri dish (12 cm × 12cm) containing uracil-, tryptophan- and methionine-free minimal agar-based solid medium (SD-Ura/Trp/Met) containing 2% (w/v) serine. Sterile filters (Thermo Fisher similar to those used for antibiograms) were placed on the agar surface, and 2 μl of individual compound from the various chemical libraries were applied to each filter. In addition, for each Petri plate, DMSO, the vehicle, was added as a negative control on the top left filter, and 2 nmol of methionine as a positive control on the bottom right filter. Plates were then incubated at 33 °C for 3 days and scanned using a Snap Scan1212 (Agfa).

Two repurposed drug libraries were screened: the Prestwick Chemical Library®(1200 drugs) and the BIOMOL’s FDA Approved Drug Library (Enzo Life Sciences, 640 drugs). In addition, the Prestwick Phytochemical library (691 green compounds, most of them being in use in Human) was also screened. The compounds were supplied in 96-well plates as 10 mM (for the two Prestwick®libraries) and 2 mg/ml (BIOLMOL®) DMSO solutions. Disulfiram was purchased from Sigma-Aldrich and resuspended in DMSO.

### Mouse model treatment with Disulfiram (DSF)

A pre-clinical protocol was designed to target cognitive defects correlated to CBS overexpression in Tg(*CBS*) mice brain (figure 4D). The selected molecule was Disulfiram (DSF), a potent inhibitor of mitochondrial aldehyde dehydrogenase (ALDH) used for the treatment of chronic alcoholism. We based our experiment on the work of Kim et al. (36) in which the DSF effect on ethanol sensitization in mice was demonstrated.

**Figure 4.**
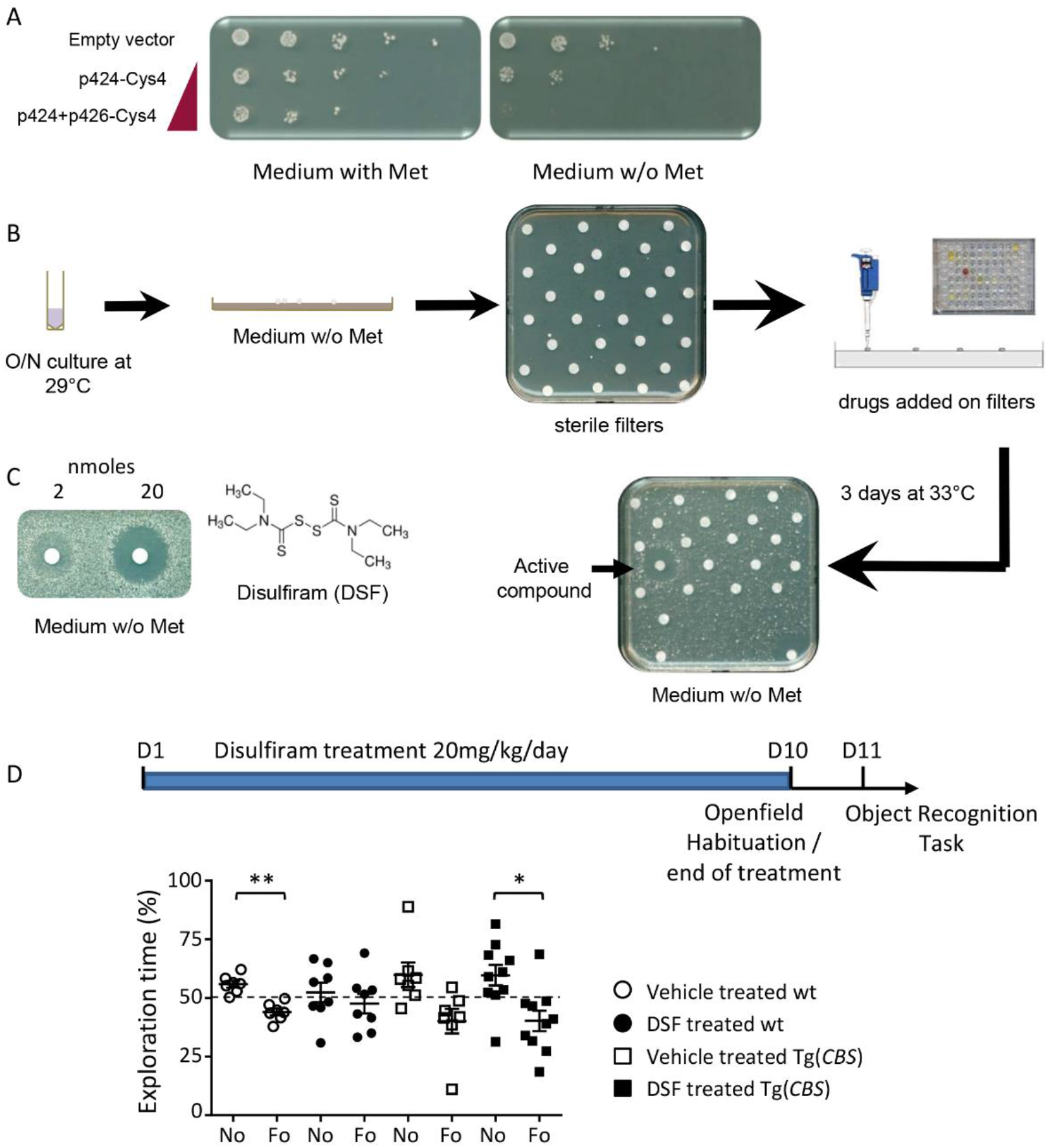
Pharmacological intervention to suppress the consequence of CBS overexpression in yeast (A, B and C) and mouse (D) Development of a yeast screening assay based on Cys4-overexpressing cells and identification of DSF as able to suppress methionine auxotrophy induced by Cys4 overexpression. The sensitivity of the strain to the absence of methionine in the medium was evidenced by serial dilutions of a yeast strain expressing different levels of Cys4 (A). For the drug screening, the yeast strain overexpressing Cys4 from both p424 & p426 multicopy 2 μ plasmids was spread on a square Petri plate containing solid agar-based methionine-free medium. DMSO was used as a negative control and added to the upper right filter and methionine, the positive control, was deposited on the bottom left filter (B). At the remaining positions, individual compounds from the chemical libraries were added, and plates were incubated for 3 d at 33 °C. The dose-dependent effect of DSF on Cys4-overexpressing cells is shown, and its molecular structure is depicted (C). Note that DSF is toxic at high concentrations (close to the filter) whereas it becomes active at sub-toxic concentrations. To test DSF in mice, a treatment was done on Tg(*CBS*) cohort starting at D1 and ending at D10 (D). Each groups received a daily dose of 10mg/kg/day of DSF for 10 days followed by an open field paradigm (D10) with the object recognition test performed on D11 (with one hour of retention time). The graph at the bottom showed the percentage of time spent on the novel versus the familiar object during the tests. The vehicule-treated wt mice were able to distinguish both objects as the DSF-treated Tg(*CBS*) animals. On the contrary non-treated transgenic animals were not able to do so and the DSF-treated wt animals were impaired in the test confirming that the drug affects CBS activity *in vivo* (Values represent means + S.E.M. *P<0.05, **P<0.01, ***P<0.001).

Behavioural studies were conducted in 12-16 week old animals; to do so, we generated 3 independent cohorts, in which we tested 4 conditions taking into account the dose of DSF (or vehicle alone) and the genotype. For the cohorts (C1 to C3), we produced respectively 5,7,3 (n=15 in total) wild type (wt) treated with vehicle, 5,3,6 (n=14 in total) transgenic for human *CBS* (Tg(*CBS*)) treated with vehicle, 7,5,3 (n=15 in total) wt treated with 10mg/kg/day of DSF, 6,6,8 (n=20 in total) Tg(*CBS*) treated with 10mg/kg/day of DSF based on the dose previously administrated in the reference publication (36). The local ethics committee, Com’Eth (n°17), approved the mouse experimental procedures, under the accreditation number APAFIS#1564-2015083114276031 with YH as the principal investigator in this study. All assessments were scored blind to genotype and animals were randomly distributed to experimental groups and treatment as recommended by the ARRIVE guidelines (37, 38). DSF was prepared at 10 mg/mL in DMSO, aliquoted and stored below -20°C. The final formulation was prepared just prior to use as a 1 mg/mL solution diluted in Cremophor EL Castor oil (BASF)/H2O ready for injection (15/75), to reach a final DMSO/Cremophor/H2O 10/15/75 (v/v/v) mix. Treated animals received a daily dose (10 days) of this formulation by intra-peritoneal injection of 10 mg/kg/day. Non-treated animals received the same formulation without DSF. On day 10 of treatment, the animal were habituated 30 min into the arena. On day 11, animals were tested in NOR paradigm to assess recognition memory after 1hour retention as described in the Open field and Object recognition task protocols (Supplementary information).

### Quantitative proteomic analysis

We collected 5 hippocampi of littermates with the 4 genotypes: wt, Dp1Yah, Tg(*Dyrk1a*)/0 and [Dp1Yah,Tg(Dyrk1a)/0] after the behavioural evaluation at the age of 25-27 weeks. Samples were reduced, alkylated and digested with LysC and trypsin at 37°C overnight. Five sets of samples with one sample from each genotypes (4 in total) were labelled with Thermo Scientific Tandem Mass isobaric tag (TMT), pooled and then analysed using an Ultimate 3000 nano-RSLC (Thermo Scientific, San Jose California) coupled in line with an Orbitrap ELITE (Thermo Scientific, San Jose California). An additional set was done comparing all the wt controls together. Briefly, peptides were separated on a C18 nano-column with a linear gradient of acetonitrile and analysed in a Top 15 HCD (Higher collision dissociation) data-dependent mass spectrometry. Data were processed by database searching using SequestHT (Thermo Fisher Scientific) with Proteome Discoverer 1.4 software (Thermo Fisher Scientific) against a mouse Swissprot database. Precursor and fragment mass tolerance were set at 7 ppm and 20 ppm respectively. Trypsin was set as enzyme, and up to 2 missed cleavages were allowed. Oxidation (M) and TMT labelled peptides in primary amino groups (+229.163 Da K and N-ter) were set as variable modification, and Carbamidomethylation (C) as fixed modification. We then compared our 5 wt samples to determine the sample closer to average score from the group, and defined it as the reference sample. All the protein quantification was done based on the reference wt sample. In total, we detected 1655 proteins filtered with false discovery rate (FDR) at 5% with a minimum of 2 peptides for a given protein detected per genotypes. We calculated the mean of the fold change for each proteins from all the samples (Dp1Yah, Tg(*Dyrk1a*) and Dp1Yah/Tg(*Dyrk1a*) compared to control. From the preliminary data, we selected 208 proteins with variability level below 40% and a fold change below 0.8 or above 1.2.

### Western Blot analysis

Ten microgram of total proteins from cortex extracts were electrophoretically separated in SDS–polyacrylamide gels (10%) and then transferred to nitrocellulose membrane (120V) during 1h30. Non-specific binding sites were blocked with 5% skim milk powder in Tween Tris buffer saline (T.T.B.S.) 1 h at room temperature. Immunoblotting was carried out with primary antibody (Supplementary table 2) incubated overnight at 4°C. The next day, we started with 3 washing baths with T.T.B.S, followed by secondary conjugated with horseradish peroxidase. The immunoreactions were visualized by ECL chemiluminescence system (Clarity_™_western ECL substrate – Bio-Rad); Epifluorescence was captured with Amersham_™_Imager 600. Bands were detected at 18, 25 and 75 kDa respectively for SNCA, SNAP25 and FUS; Signals were quantified with ImageJ.

## RESULTS

### Three copies of *Cbs* are necessary to induce cognitive impairments in the Dp1Yah mice

In order to challenge the hypothesis that three copies of *Cbs* are necessary to induce behavioural deficits in the Dp1Yah mice, we combined the Dp1Yah mice with the *Cbs^tm1Unc/+^* knock-out model (29) and we compared the Dp1Yah with Dp1Yah/*Cbs^tm1Unc^* (in which only two copy of *Cbs* are functional), wild type (wt) and *Cbs^tm1Unc/+^* heterozygote controls. In the open field test, most of the genotypes displayed similar exploratory behaviour, except for the Dp1Yah/*Cbs* mice that travelled more distance in the open field arena with a higher speed (Figure 1A left panel; On way ANOVA on distance, post hoc Tukey Test: Dp1Yah vs Dp1Yah/*Cbs^+/tm1Unc^* p=0.002; Figure 1A right panel; On way ANOVA on speed, post hoc Tukey Test: wt vs Dp1Yah/*Cbs^+/tm1Unc^* p=0.004; *Cbs^+/tm1Unc^* vs Dp1Yah/*Cbs^+/tm1Unc^* p=0.05; Dp1Yah vs Dp1Yah/*Cbs^+/tm1Unc^* p=0.007). Similarly when the mice performed the Y maze, we confirmed the increased activity with a higher number of arm entries for the Dp1Yah/*Cbs^tm1Unc^* compared to the other genotypes (Figure 1B; Kruskal-Wallis One way ANOVA on Ranks – genotypes, post hoc Dunn’s method: wt vs Dp1Yah/*Cbs^+/tm1Unc^* p<0.05; Dp1Yah vs Dp1Yah/*Cbs^+/tm1Unc^* p<0.05) but no impact on spontaneous alternation (One way ANOVA, F(3,87)=2.486 p=0.066). To determine if motor activity was altered in the Dp1Yah/*Cbs^tm1Unc^* model, we used the rotarod test. After the first day of training we did not find any change in the maximum speed reached before falling for all tested genotypes (Figure 1C; Speed: repeated measures ANOVA variable « genotype » and « day », F(3;110)=1.816 p=0.155). Nevertheless, we observed a decrease in the locomotor learning in the Dp1Yah mice comparing to the next following days of training which was rescued in the Dp1Yah/*Cbs^tm1Unc^* mutant (Figure 1C; Speed: repeated measures 2 way ANOVA variable « genotype » and « day », F(2;165)=17.171 p<0.001 post hoc Tuckey method wt «day1 vs day3» p=0.002; *Cbs^tm1Unc^*^/+^ «day1 vs day3» <0,001; Dp1Yah «day1 vs day3» p=0.238; Dp1Yah/*Cbs^tm1Unc^* «day1 vs day3» p=0.017). During the test phase, we found that the Dp1Yah individuals showed a weaker performance compared to *Cbs^tm1Unc^*^/+^ and Dp1Yah/*Cbs^tm1Unc^* (ANOVA, variable « speed » and « genotype » F(3;385)=5.544 p<0.001 post hoc Tuckey method; «wt vs Dp1Yah» p=0.099; «*Cbs^tm1Unc^*^/+^ vs Dp1Yah» p=0.001; «Dp1Yah vs Dp1Yah/*Cbs^tm1Unc^*» p=0.01).

**Figure 1.**
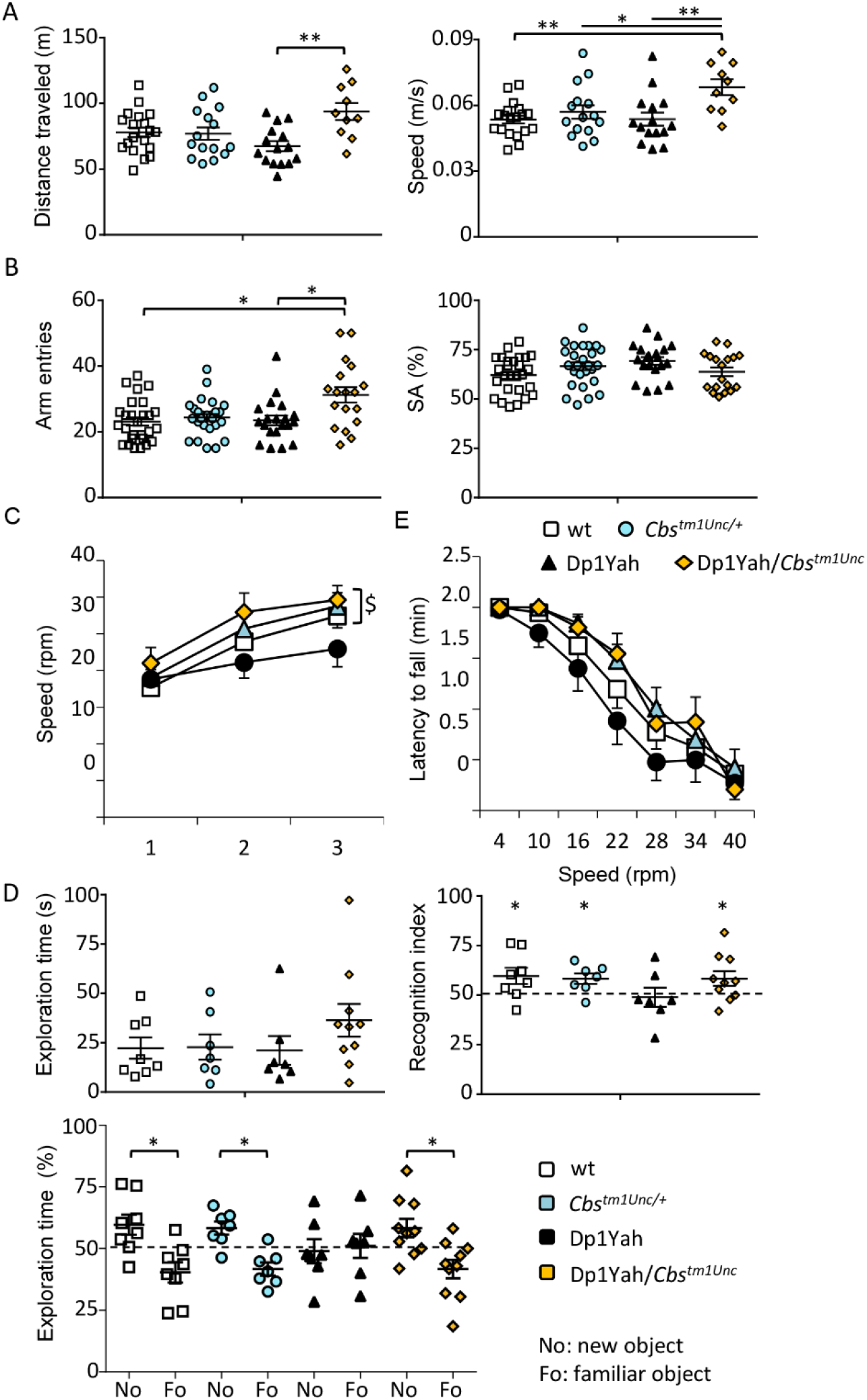
The Dp1Yah phenotypes are dependent on *Cbs* dosage. Dp1Yah trisomic mice (n=23) were compared with Dp1Yah carrying a KO of *Cbs* (Dp1Yah/*Cbs^tm1Unc^*, n=21), *Cbs^tm1Unc/+^*(n=23) and wt littermates (n=29). Animals were analysed for the open field (A), the Y maze (B) and the novel object recognition (D) in two independent cohorts; the rotarod (C) was assessed on one cohort with wt (n=18) *Cbs^tm1Unc/+^* (n*=*15), Dp1Yah (n=15) and Dp1Yah/*Cbs^tm1Unc^* (n=10) littermates. (A) Distance travelled and medium speed during the 30min of the test were increased in the *Dp1Yah/Cbs^tm1Unc^* compared to the wild type genotype. (B) Increased exploration activity was confirmed for the *Dp1Yah/Cbs^tm1Unc^* mice compared to control littermates in the Y maze while spontaneous alternation was not affected. (C) During the training session (left panel), the Dp1Yah mice were not able to improve their performance on the rotarod by increasing the maximum of speed before they fall from the rod compared to the other genotype. Nevertheless no change was observed between individuals with the four genotypes during the test phase (right panel). (D) The exploration time in the first session of the novel object recognition (left upper panel) was not statistically different in the four genotypes but during the recognition phase, after 10 min of retention, the recognition index (right upper panel; time spent on the new object / total time of exploration) was clearly lower in Dp1Yah mice as compared to the other genotypes and not statistically different from chance (50%). Accordingly the exploration time (left lower panel) spent by the *Dp1Yah/Cbs^tm1Unc^* mice to explore the object showed that they were able to differentiate the novel (No) versus the familiar (Fo) object while the Dp1Yah were not. Data are represented as one point per individual tested and the mean of the group. (Values represent means + S.E.M. *P<0.05, **P<0.01, ***P<0.001).

Then we tested the object memory. No difference was observed during the exploration of the familiar object in the presentation phase of the test (Figure 1D top left panel). However, during the discrimination phase, after 1h of retention, the Dp1Yah mutant mice were not able to differentiate the familiar versus the novel object whereas the wt, *Cbs^tm1Unc/+^* and the Dp1Yah/*Cbs^tm1Unc^* spent significantly more time on the new object compared to the familiar one (Figure 1D, left bottom panel; two ways ANOVA, variables “genotype” and “objects”: F(3;56)= 2.86 with p=0.045; post hoc Tuckey method wt “fam vs new” q= 4.885 and p= 0.001; *Cbs^tm1Unc/+^* q= 3.913 and p= 0.008; Dp1Yah, q= 0,503 and p= 0.724; Dp1Yah/*Cbs^tm1Unc/+^* q= 4.715 and p= 0.002). Accordingly, the recognition index showed that the restoration of two functional copies of *Cbs* in the Dp1Yah mice rescued memory performance in object recognition (Figure 1D right panel; One sample t-test: wt p=0.05; *Cbs^tm1Unc/+^* p= 0.01; Dp1Yah p=0.82; Dp1Yah/*Cbs^tm1Unc/+^* p=0.05).

Overall this set of experiments demonstrated that 3 copies of *Cbs* were necessary for inducing the Dp1Yah phenotypes in novel object recognition. In addition rescuing *Cbs* dosage induced a slight hyperactive phenotype during the exploration of a new environment and restored performance in the rotarod activity. Interestingly, returning back to wt level of expression of *Cbs* in the *Abcg1-Cbs* region enables another trisomic gene from this region to impact on the exploratory behaviour of the mouse

### The sole overexpression of a human *CBS* transgene impacts the object recognition and the locomotor activity

We used the Tg(*CBS*), a PAC transgenic line encompassing a 60kb fragment with the human *CBS* locus (30) to analyse the impact of the sole increase of *Cbs* dosage on behaviour and cognition. As shown in figure 2A, no difference in locomotor activity was observed during the exploration of a new environment in the open field test between wt and transgenic littermates (Student t-test distance: wt vs Tg(*CBS*)/0 p=0.925; speed wt vs Tg(*CBS*)/0 p=0.925). However we found higher circadian activity for isolated individuals (Figure 2C; student t-test wt vs Tg(*CBS*) p<0.001) which results from an increased locomotor activity during the habituation and the dark phase (Figure 2B). In the Y maze (Figures 2D-E), no difference was detected for the number of arm entries and the spontaneous alternation. In the novel object recognition test, (Figures 2F-H) the Tg(*CBS*)/0 animals spent more time sniffing the two identical objects during the presentation phase than their control littermates (Figure 2F; Student t-test wt vs Tg(*CBS*)/0 p=0.05) but were impaired in object recognition as shown by the absence of discrimination between novel and familiar objects for the transgenic mice (Figure 2G: Student paired t-test wt “Fo vs No” p= 0.008; Tg(*CBS*) “Fo vs No” p=0.174) resulting in a recognition index (time on the new object / total time) not significantly different from the 50% chance level, (Figure 2H: one sample t test, significant difference from 50%, wt p= 0.008; Tg(*CBS*)/0 p= 0.174). Consequently we demonstrated that CBS overexpression is sufficient to induce deficit in novel object recognition memory and decreased locomotor activity during dark phase while having no effect during the light phase.

**Figure 2.**
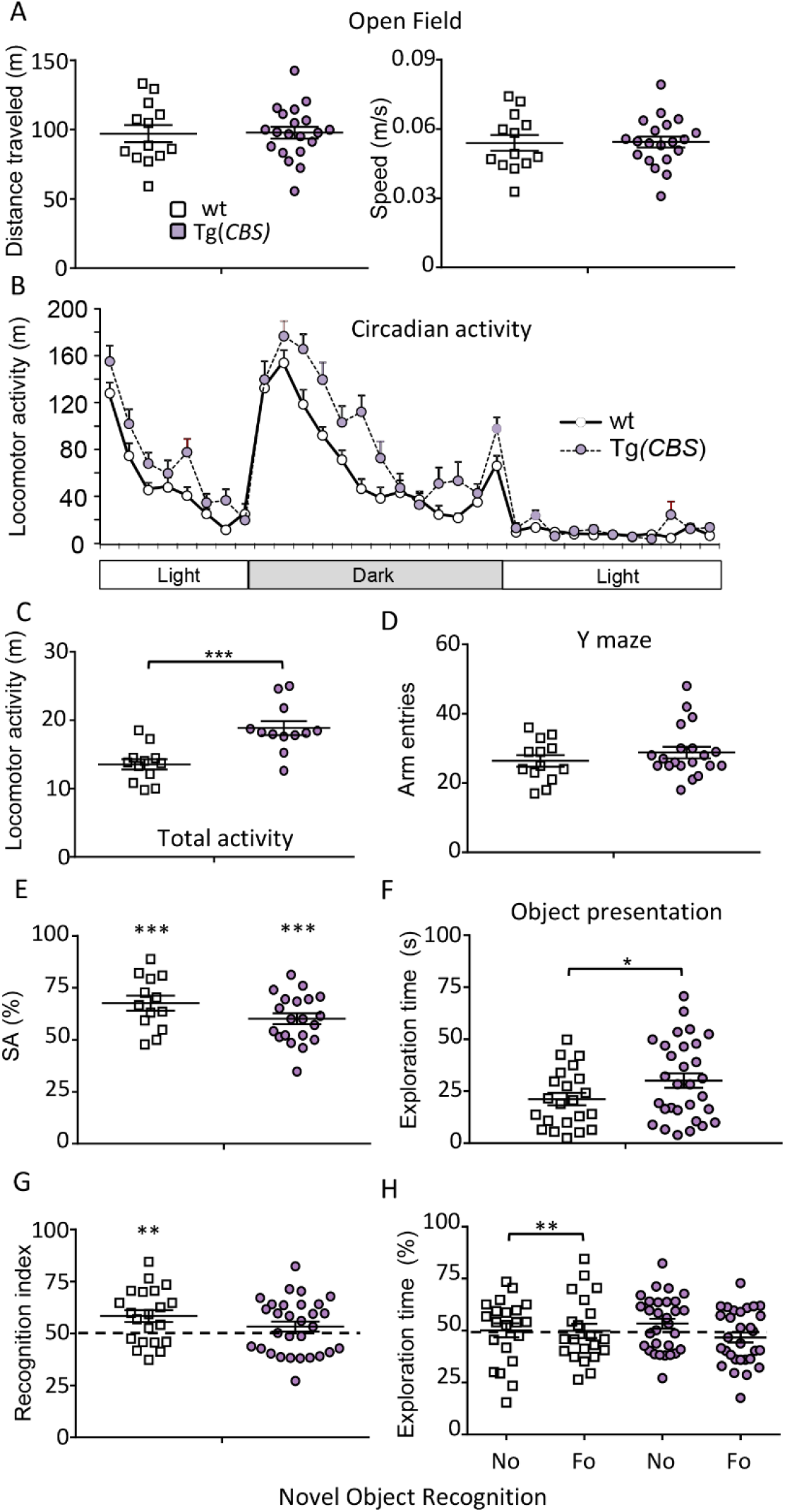
Transgenic mice overexpressing human CBS display DS-related behaviour phenotypes. Wt(n=13) and Tg(CBS)/0 littermates (n=17), hemizygotes for a human PAC containing the *CBS* gene, were tested for open field (A), circadian actimetry (B,C), Y maze (D-E) and novel object recognition (F,G and H). No phenotype was found in the Tg during the exploration of a new environment in the open field in the total distance travelled (left) and the speed (right) but increased activity was observed during home cage monitoring over a light-dark-light cycle (B) with an increase of the distance travelled (C). In the Y maze (E), Tg(*CBS*)/0 animals displayed altered spontaneous alternation with no change in the number of arm entries (D). In the novel object recognition (F), Tg(*CBS*)/0 mice displayed similar exploration activity compared to wt littermates but they do not discriminate the novel versus the familiar object when looking at the discrimination index (G) and the percentage of exploration time for both objects (H). (Values represent means + S.E.M. *P<0.05, **P<0.01, ***P<0.001).

### Cbs overexpression in hippocampal and cortical neurons induces behavioural defects similar to Dp1Yah

We checked if we could induce the cognitive deficits observed in DS mouse models by overexpressing *Cbs* mostly in the hippocampal and cortical neurons involved in learning and memory. Hence we engineered the Tg(*Prp-gfp-CBS*) mouse strain in which the human CBS cDNA can be expressed from the Prion promoter after the excision of the gfp cassette flanked by loxP sites (Figure 3A) and selected one Tg(*Prp-gfp-CBS*) line with a pattern of expression in the anterior part of the adult brain (Figure 3B). We chose the Tg(*Camk2a-cre*) (31), to direct the cre expression in the cortical and hippocampal glutamatergic neurons and we verified the expression of the human *CBS* in different brain regions of the double transgenic (Tg(*Prp-gfp-CBS*)/0;Tg(*Camk2a-cre*)/0). As expected we found expression levels comparable to the endogenous murine *Cbs* gene in cerebellum while human CBS was overexpressed in the hippocampus and the cortex (Figure 3C). Littermate animals carrying wt, the two single transgenic constructs and the two transgenes were produced and tested for object recognition. During the test, the control groups, namely wt, Tg(*Prp-gfp-CBS*)/0 and Tg(*Camk2a-cre*)/0, spent more time on the new object (No) than the familiar one (Fo) as expected, while the double transgenic individuals were not able to differentiate the new object from the familiar one as shown by the recognition index or the percentage of exploration time (Figure 3D; Recognition index: One sample t-test: wt p=0.03; Tg(*Camk2a-cre*)/0 p=0.03; Tg(*Prp-gfp-CBS*)/0 p=0.001; (Tg(*Prp-gfp-CBS*)/0; Tg(*Camk2a-cre)/0)* p=0.90; exploration time; two ways ANOVA, variables “genotype” and “objects”: F(3; 76)= 8.59 with p<0.001; post hoc Tuckey method wt «No vs Fo» p<0,001; Tg(*Camk2a-cre*)/0 «No vs Fo» p=0.001 and Tg(*Prp-gfp-CBS)/0* «No vs Fo» p<0.001; (Tg(*Prp-gfp-CBS*)/0; Tg(*Camk2a-cre)/0*)) «No vs Fo» p=0,861).

Measurements of the travelled distance in the open field and number of visited arms in the Y maze revealed hyperactivity of the Tg(*Camk2cre)*/0 carrier groups (Figures 3E-F; Openfield: One way ANOVA F(3,49)=4.80 p=0.005; post hoc Holm-Sidak «wt vs Tg(*Camk2*-*Cre*)/0» unadjusted p=0.002; «Tg(*Prp-gfp-CBS*)/0 vs Tg(*Camk2*-*Cre*)/0» p=0.003) - Y maze: One way ANOVA F(3,46)=6.04 p=0.001; post hoc Holm-Sidak «wt vs Tg(*Camk2*-*Cre*)/0» p=0.04; «Tg(*Prp-gfp-CBS*)/0 vs Tg(*Camk2*-*Cre*)/0» p=0.009; Tg(*Prp-gfp-CBS*)/0 vs Tg(*Prp-gfp-CBS*)/0;Tg(*Camk2a-cre)/0*p=0.04). Like for the Dp1Yah and Tg(*CBS*) animals, we did not found any alteration in the spontaneous alternation in the Y maze test (One way ANOVA: F(3,43)=0.691 p=0.563). All the mice, whatever their genotype, performed equally well during the training session of the rotarod (Figure 3G) (training: repeated measures ANOVA, variables « genotype » and « day », F(3;90)=2.011 p=0.126; test: repeated measures ANOVA, variables « genotype » and « day », F(2;90)=44.783 p<0.001) as well as during the test session with increasing speed (Repeated measures ANOVA, variables « genotype » and « speed », F(18;322)=0.631 p=0.875). Thus, as expected from the role of the cerebellum in locomotor coordination, the overdose of CBS restricted to cortical and hippocampal neurons did not interfere with the locomotor activity.

Hence, overexpression of CBS is necessary and sufficient to induce object memory defect in a 1h retention test with limited impact on other phenotypes. As such, *CBS* is a new gene whose overdosage alters cognition in DS mouse models and as a consequence is likely to contribute to DS phenotypes.

### Identification of drugs that suppress the effects of Cys4/CBS overexpression both in yeast and mouse

A few studies have reported the identification of CBS inhibitors (39-44) but most of them were based on *in vitro* assays using a recombinant CBS enzyme as a drug target and led to the isolation of inhibitors with relatively low potency and limited selectivity, hence leading to the idea that CBS may be an undruggable enzyme. Therefore we oriented toward an *in cellulo* phenotype-based assay that would allow screening drugs that interfere with the phenotypical consequences of CBS overexpression and thereby that do not necessarily directly target the CBS enzyme. The budding yeast *Saccharomyces cerevisiae* contains a functional homolog of CBS and has been shown to be a relevant system to model pathophysiological mechanisms involved in a number of human disorders and to perform chemobiological approaches that aim at identifying both drugs and new therapeutic targets (45-51). We thus decided to create a yeast model in which the phenotypical consequences of CBS overexpression may be easily and conveniently monitored in order to get a potential *in cellulo* high throughput drug screening procedure. We reasoned that if we overexpressed CBS at a sufficient level, this should lead to a decreased intracellular level of methionine, similarly to what was shown in patients, and therefore that yeast cells would become methionine auxotroph and thereby unable to grow on methionine-free minimal media. As the human CBS protein is not very stable in yeast cells and therefore cannot be expressed at high levels (52), we decided to overexpress Cys4p, the CBS homolog in *S. cerevisiae*. Cys4p presents the same domains and domain organization than CBS apart from the N-terminal heme-binding domain which is absent in the yeast protein (53). To get a degree of methionine auxotrophy sufficient to allow an efficient screening, we expressed Cys4 from the strong constitutive *GPD* promoter from two different high copy number 2 μ vectors (each present at ~50 copies per cell) and supplement the growth medium with serine, which is one of the Cys4p/CBS substrates that could otherwise become limiting upon Cys4 overexpression (Figure 4A).

Using this model, we tested ≈ 2200 compounds from 3 different chemical libraries consisting mainly of repurposed drugs for their ability to suppress the methionine auxotrophy induced by Cys4p overexpression. We exploited a similar principle as a yeast-based screening setup previously (46, 47, 49, 54). Briefly, we spread, on a solid agar-based methionine-free minimal medium, yeast cells overexpressing Cys4. Then we put filters on the agar surface and add different drugs from chemical libraries on each filters. After 3 days of incubation at 33°c, active compounds were identified by a halo of restored/enhanced growth around the filter on which they were loaded (Figure 4B). The advantage of this method is that, in one simple experiment, it allows numerous compounds to be tested across a large range of concentrations due to the diffusion of the molecule in the medium surrounding the filter onto which it was deposited. This design drastically improves the sensitivity of the screen because the screened compounds can be toxic at high concentrations whereas being active at subtoxic concentrations. We identified four different compounds, among which disulfiram (DSF, Figure 4C).

Next we tested if DSF was able to restore the object recognition of the mouse model overexpressing human *CBS*. Three independent cohorts of Tg(*CBS*) and control littermates were treated with DSF (10mg/kg/day) for 10 days before being tested for the novel object recognition. As shown in figure 4D, DSF-treated transgenic animals were restored in the novel object recognition paradigm whereas non treated mutant animals were still not able to discriminate the new versus the familiar object. Interestingly the wt treated individuals were no more able to perform the discrimination while the vehicle treated controls were able to do so (Student paired t-test: vehicle treated wt «No vs Fo» p=0,006; DSF treated wt «No vs Fo» p=0.11 and vehicle treated Tg(CBS) «No vs Fo» p=0.59; DSF treated Tg(*CBS*) «No vs Fo» p=0,05). This goes in line with the fact that loss-of-function mutations in CBS also leads to cognitive defects as observed in homocystinuria patients. Hence, this latter result confirm that DSF does affect CBS activity, directly or indirectly. Altogether these results confirm that the phenotypical consequences of the overexpression of CBS could be targeted by drugs to restore some of the cognitive performance altered in DS models. They also emphasize that the inhibition of CBS, direct or indirect, should be mild and only partial as a strong inhibition may be detrimental as illustrated by the cognitive dysfunction observed in homocystinuria and here in wt mice treated with DSF.

### Epistatic interaction between *Dyrk1a* and the *Abcg1-Cbs* region drives recognition memory in DS mouse models

*Dyrk1a* is a major driver gene of DS cognitive defects (55) and a decrease in *Cbs* dosage is known to change the expression of *Dyrk1a* in brain and other organs (56-58). Thus in order to test the functional interaction of *Cbs* and *Dyrk1a* overdosage, we combined the Dp1Yah with the Tg(*Dyrk1a*) mouse model, with *Dyrk1a* mRNA expression ratio around 1.5 compared to control littermate (32). Tg(*Dyrk1a*) mice present increased spontaneous activity compared to wt in the Open field test. This hyperactivity was also observed in the double transgenic Dp1Yah/Tg(*Dyrk1a*) while it was absent from Dp1Yah animals (Figure 5A; Student t test wt vs Dp1Yah p=0,460; wt vs Tg(*Dyrk1a*) p=0.002 and wt vs Dp1Yah/Tg(*Dyrk1a*) p=0.006; Tg(*Dyrk1a*) vs Dp1Yah/Tg(*Dyrk1a*) p=0,200). Hyperactivity was confirmed in the Y-maze, with both Tg(*Dyrk1a*) and Dp1Yah/Tg(*Dyrk1a*) having more arms visits than the controls and Dp1Yah (Figure 5B; Student t test wt vs Dp1Yah p=0,800; wt vs Tg(*Dyrk1a*) p=0.005 and wt vs Dp1Yah/Tg(*Dyrk1a*) p=0.005; Tg(*Dyrk1a*) vs Dp1Yah/Tg(*Dyrk1a*) p=0,881). The working memory defect observed in the Y maze for Tg(*Dyrk1a*) mice was not rescued in Dp1Yah/Tg(*Dyrk1a*) double transgenics (Figure 5B; One way ANOVA F(3,48)=4.14 p=0.011; post hoc Tukey method wt vs Tg(*Dyrk1a*) p=0.042; wt vs Dp1Yah/Tg(*Dyrk1a*) p=0,019 and Tg(*Dyrk1a*) vs Dp1Yah/Tg(*Dyrk1a)* p=0.203). Then, we tested the Novel Object Recognition memory after 1h of retention (Figure 5C). As expected, the 2 single mutants were impaired (Two ways ANOVA, variables “genotype” and “objects”: F(3;70)=7.09 with p<0.001, post hoc Tukey Test: Dp1Yah “fam vs new” q=1.333 and p=0.349; Tg*(Dyr1a)* q=1.732 and p=0.225 - Recognition Index: One sample t-test mean vs 50%: Dp1Yah p=0.253; Tg(*Dyrk1a*) p=497) but the double transgenic mice Dp1Yah/Tg(*Dyrk1a*) were able to discriminate the novel object as wt littermates (Two ways ANOVA, variables “genotype” and “objects”: F(3;70)=7.09 with p<0.001, post hoc Tukey Test: wt “fam vs new” q=4.543 and p=0.002; Dp1Yah/Tg*(Dyr1a)* q=5.289 and p<0.001 - Recognition Index: One sample t-test: wt p=0.048; Dp1Yah/Tg(*Dyrk1a*) p=0.011), suggesting that the effects of *Dyrk1a* overexpression are compensated by 3 copies of the *Abcg1-Cbs* region.

**Figure 5.**
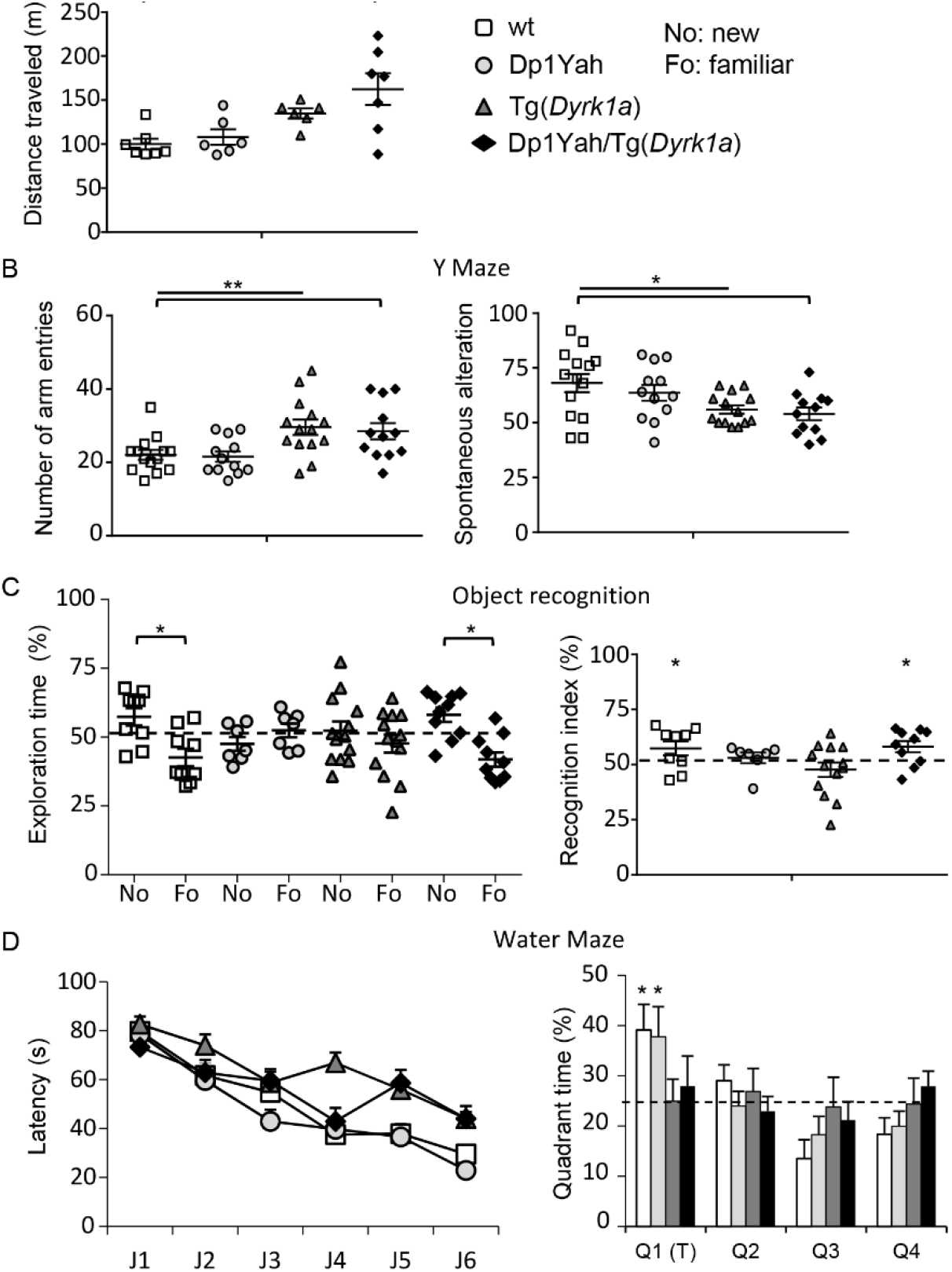
CBS and DYRK1A overdosages interact for controlling behaviour and cognition. Behavioural and cognitive analysis of transgenic animals overexpressing *Cbs* and *Dyrk1a* (14 wt, 15 Tg(*Dyrk1a*), 13 Dp1Yah and 13 Dp1Yah/Tg(*Dyrk1a*)) mutant mice in the open field (A), the Y maze (B), the object recognition (C) and the Morris water maze (D). Increased activity in the open field (A) and in the number of arm entries in the Y maze (B) were found in the Tg(*Dyrk1a*) and Dp1Yah/Tg(*Dyrk1a*) animals with also reduced spontaneous alternation in the Y maze (B). Both the Dp1Yah and Dp1Yah/Tg(*Dyrk1a*) mutant mice were impaired in object recognition (C) but the double mutant animals showed restored object discrimination similar to wt littermates. The Tg(*Dyrk1a*) and Dp1Yah/Tg(*Dyrk1a*) animals displayed delayed learning in the Morris water maze with no memory of the platform location in the probe test compared to Dp1Yah and wt littermates (D). (Values represent means + S.E.M. *P<0.05, **P<0.01, ***P<0.001)

Lastly we checked the learning and spatial memories using the Morris Water Maze task, followed by a probe test 24h after the learning period (Figure 5D). Even if all the groups increased their performance during the learning phase for reaching the platform after 6 days of training (J1-J6), wt and Dp1Yah mice found the platform with lower latency than the Tg(*Dyrk1a*) and Dp1Yah/Tg(*Dyrk1a*) (Two ways ANOVA variable genotype, F(3;280)=14.80 p<0.001; post hoc Tuckey test: wt vs Tg(*Dyrk1a*) q=6.160 with p<0,001; wt vs Dp1Yah/Tg(*Dyrk1a*) q=4.752 with p=0.004 – Dp1Yah vs Tg(*Dyrk1a*) q=8.103 with p<0,001; Dp1Yah vs Dp1Yah/Tg(*Dyrk1a*) q=6.641 with p<0,001). During the probe test, 24h after the learning phase, controls and Dp1Yah animals were searching most of their time in the platform quadrant (T), whereas Tg(*Dyrk1a*) and double transgenic mice searched randomly across the entire space (One sample t-test vs 50% mean: wt p=0.02; Dp1Yah p=0.05; Tg(*Dyrk1a*) p=0.99 and Dp1Yah/Tg(*Dyrk1a*) p=0.57). Hence, overexpressing Cbs and Dyrk1a does not rescue the Dyrk1a-dosage dependent working and spatial memory deficits observed in the Y maze and the Morris water maze respectively neither the hyperactivity observed in the open-field, but rescued the object recognition impairment in the NOR.

### Proteomics unravels complex intermingled proteomic changes influenced by DYRK1A overexpression and by Dp1Yah trisomic genes

In order to unravel the impact of CBS and DYRK1A on cellular mechanism within the hippocampus that could lead to the memory phenotype observed in the novel object recognition (NOR) test, we profiled the proteome in the hippocampi isolated from Dp1Yah, Tg(*Dyrk1a*) and double (Dp1Yah,Tg(*Dyrk1a*)) animals, and compared them to the wt control littermates. We collected the samples after the behavioral evaluation and performed a Tandem Mass Tag labeling (Thermo Scientific, Illkirch) followed by LC-MS/MS orbitrap analysis. We were able to detect 1655 proteins of which 546 were detected in all the 3 genotypes with a variability below 40% (Supplementary table 3), and among which 338 proteins were expressed at the same level as control ones. A total of 208 proteins were found differentially expressed with levels of expression above 1.2 (206) or below 0.8 (2) in Dp1Yah, Tg(*Dyrk1a*) and double mutant mice (Figure 6A). Nine proteins were upregulated in all 3 genotypes: the RIKEN cDNA 6430548M08 gene product (6430548M08RIK), Actin related protein 2/3 complex, subunit 1A (ARPC1A), Bridging Integrator 1 (BIN1), the Family with sequence similarity 213, member A (Fam213a), Glyoxalase 1 (GLO1), Importin 5 (LPO5), NADH dehydrogenase (ubiquinone) Fe-S protein 1 (NDUFS1), Prostaglandin reductase 2 (PTGR2) and Synaptosomal-associated protein 25 (SNAP25). Toppcluster analysis of the protein content unraveled a general common network with interacting proteins modified by the 3 genetic conditions (Figure 6B-C). Functional analysis using gene ontology highlighted several cellular components affected in the 3 genotypes including synaptic particles, neuron projection, presynapse/synapse, axon, myelin sheath and different types of vesicles (Supplementary table 4). Cell/neuron projection development, morphogenesis, and differentiation, as well as secretion, synaptic and anterograde trans-synaptic signaling were affected in Dp1Yah while aldehyde catabolic processes and regulation of anatomical structure size were modified in Tg(*Dyrk1a*). Interestingly all these biological process were not be disturbed in double transgenic animals. Likewise molecular functions controlling ubiquitin protein ligase, calcium ion binding and dicarboxylic acid transmembrane transporter activity in Dp1Yah, or cytoskeletal protein and myosin binding in Tg(*Dyrk1a*) were restored (Dp1Yah,Tg(*Dyrk1a*)). On the contrary oxidoreductase activity was newly modified in the double transgenic hippocampi.

**Figure 6.**
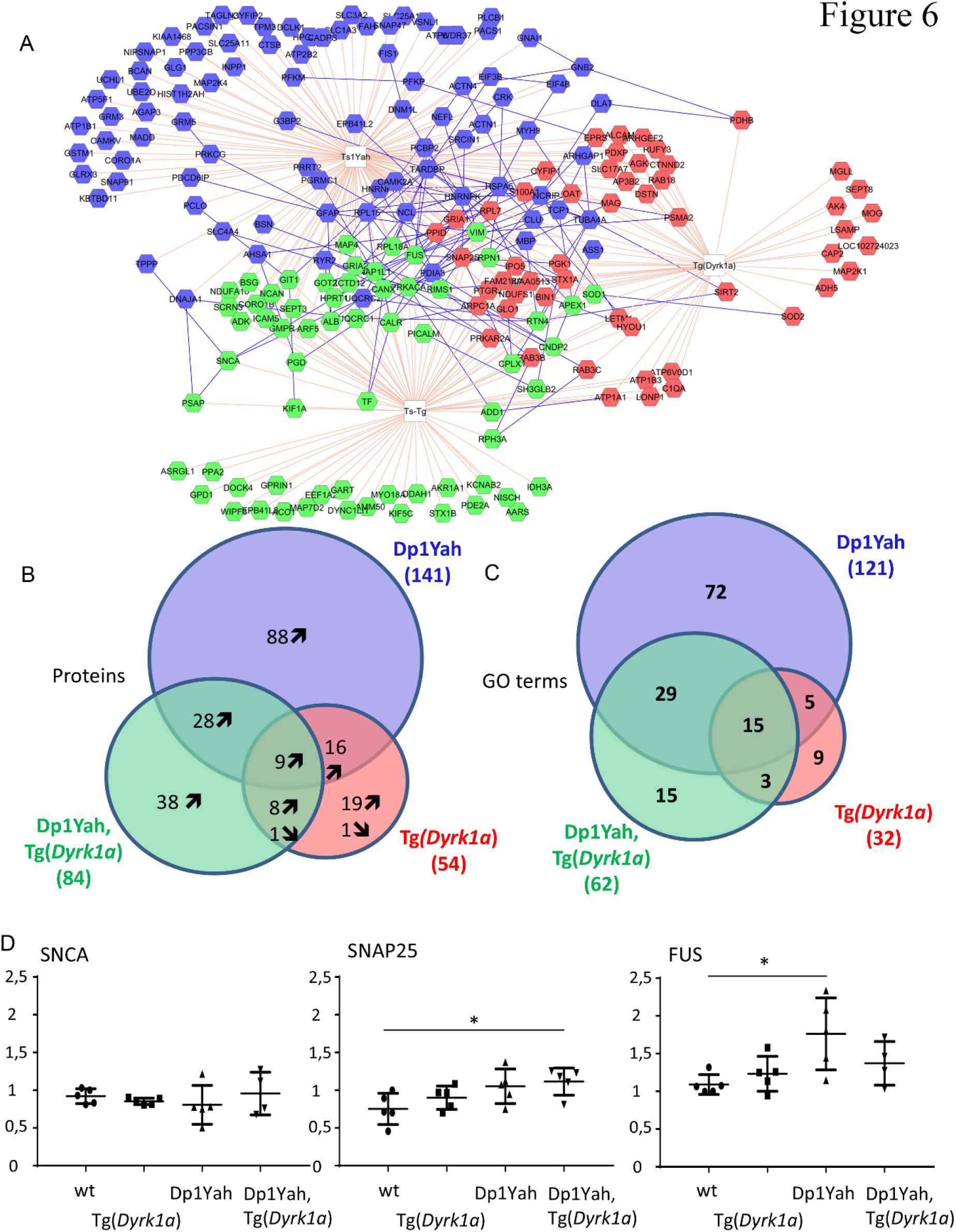
Pattern of protein expression is disrupted upon changes in DYRK1A and CBS dosage. (A) Analyzing the 1655 proteins detected in the Orbitrap ELITE experiment, we extracted from Proteome Discoverer 1.4 © a list of 208 proteins dysregulated in our different sample conditions. The association between proteins, pathways and genotype is summarized in two Venn diagrams (B-C). We deduced that the trisomic alleles induced most of the perturbations; moreover, the combination of increased DYRK1A and trisomic condition leaded to new dysregulations. (D) Western blot validation of 3 protein candidates SNCA, SNAP25 and FUS. SNAP25 expression is increased in samples overexpressing DYRK1A. More interestingly, FUS was found significantly upregulated in Dp1Yah - plots represent every sample values normalized with -actin level). (Values represent means + S.E.M. *P<0.05, **P<0.01, ***P<0.001)

We selected three proteins with different proteomic profiles in hippocampi and studied their expression in another brain region, the cerebral cortex, using western blot analysis: The alpha synuclein (SNCA), the Fused in sarcoma (FUS) that are associated with neurodegenerative disease (59-62) and the synaptosomal-associated protein 25 (SNAP25), a component of the SNARE complex involved in calcium-triggered exocytosis (63-65). As shown in figure 5D, levels of SNCA were similar to wt level. We did observe increased amount of this protein in the (Dp1Yah,Tg(*Dyrk1a*)) animals contrary to what was observed in the proteome analysis. The presynaptic SNAP25 protein was significantly up-regulated in cortical regions of the (Dp1Yah,Tg(*Dyrk1a*)) animals and to a lesser extent in the Dp1Yah and Tg(*Dyrk1a*) ones (student t-test wt versus Tg(*Dyrk1a*) p=0,233; wt versus Dp1Yah p=0,06; wt versus D1Yah/Tg(Dyrk1a) p=0,02). Hence, in the proteomic approach, we also observed the increase previously detected in the hippocampus of those three transgenic lines. The RNA-binding protein FUS was found overexpressed in the Dp1Yah brains and to a lesser extent in the (Dp1Yah,Tg(*Dyrk1a*)) ones, similarly to what was observed in the proteomic analysis (student t-test Dp1Yah compared to wt p=0,02 and D1Yah/Tg(Dyrk1a) compared to wt p=0,09).

## DISCUSSION

In this report we demonstrated that the genetic overdosage of *Cbs* is necessary and sufficient to induce defective novel object recognition in 3 different types of DS models. CBS overdosage is certainly the main driver of the learning and memory phenotypes detected previously in DS models for the Mmu17 region (22, 33) but we cannot rule out the possibility that one or more other gene(s) contribute with *Cbs* to the phenotype. Previous analysis of CBS overdosage with the same transgenic line Tg(*CBS*) on the FVB/N genetic background showed no change in fear learning task and locomotor activity but increased LTP-dependent synaptic plasticity (66); a phenomenon also detected *in vitro* and *in vivo* in other DS models where *Cbs* is trisomic in the C57BL/6J genetic background (22, 33). Nevertheless no positive effect on cognition is associated with increase CBS dosage as previously proposed by Régnier et al. (66). Instead the overdosage of CBS always impairs the hippocampal-dependent novel object recognition test suggesting that increased synaptic plasticity found in *Cbs* trisomic models may alter synaptic functions. Increased synaptic plasticity could occur via increased H2S as it has been shown that H2S facilitates LTP by stimulating the post-synaptic NMDA receptors (67, 68). Moreover, a role of H2S has been foreseen in calcium homeostasis regulation which is also crucial for neuronal synaptic plasticity (69).

DSF was isolated from a drug screening performed in yeast cells overexpressing CBS homolog Cys4p and looking for drugs counteracting its effect on methionine auxotrophy. Although DSF has been first identified as an inhibitor of mitochondrial aldehyde dehydrogenase (ALDH) (70), it is a relatively nontoxic substance, which has been on the market for more than 40 years to support the treatment of chronic alcoholism by producing an acute sensitivity to ethanol, thanks to its ability to inhibit aldehyde dehydrogenases, thus leading to an accumulation of acetaldehyde in blood when alcohol is ingested. As acetaldehyde is responsible for many of the unpleasant effects that follow ingestion of large quantities of alcohol (“hangover”), DSF treatment discourages the patients to sustain a regular alcohol consumption by exacerbating and accelerating its unpleasant side effects. Our preliminary data about the mechanism of action of DSF suggest that this molecule may not directly inhibit CBS enzymatic activity but probably rather acts on the cellular consequences of CBS overexpression. The assay used for the screening, in principle, leads to the isolation of drugs acting both directly or not on CBS/Cys4. This latter point is of importance given that CBS may not be a druggable target enzyme. And indeed, at present, we do not know if the DSF is acting directly or indirectly on CBS but we must assume the function altered by CBS overdosage, whatever it is, is conserved and similarly sensitive to DSF treatment in both yeast and mouse. Of note, upon absorption DSF is rapidly reduced to diethyldithiocarbamate (DDC), which then reacts with thiol groups. Both DSF and DCC are potent copper chelators, thereby possibly affecting the activity of copper-dependent enzymes such as monooxygenases, the Cu-Zn superoxide dismutase, amine oxidase, ADN methyltransferases and cytochrome oxidase. As a result, DSF has been shown to affect various cellular processes such as cocaine metabolism and catecholamine synthesis, and proteasome inhibition, and is thus under study for multiple clinical applications that include struggle against alcohol addiction, cancer chemotherapy, treatment of copper-related disorders and anti-viral treatment for hepatitis C and Human Immunodeficiency Virus (71). Here, we describe a new possible clinical application of DSF in DS cognition through its effect on CBS overexpression. CBS clearly represents a new relevant therapeutic target for improving DS cognition and DSF, as such, opens new therapeutic avenues in DS patients.

We also demonstrated that *CBS* interacts genetically with *Dyrk1a*, a well-known therapeutic target for DS. Mutual relationships between DYRK1A and CBS were shown previously, with decreased DYRK1A protein observed in the liver (Hamelet et al. 2009) and increased expression observed in the brain of *Cbs^+/-^* mice (Planque et al. 2013), while overexpression (or under-expression) of DYRK1A induce accumulation (or reduction) of CBS expression in the liver (72). In order to explore the genetic interactions between DYRK1A and CBS, we overexpressed *Dyrk1a* in the Dp1Yah context by combining the Tg(*Dyrk1a*) and the Dp1Yah mice. Surprisingly, this experiment restored the object recognition deficit observed in the Dp1Yah mouse model but neither the increased locomotor activity in the open-field or the Y maze, nor the working and spatial memory deficits. Thus the compensation is restricted to a specific cognitive function, recognition memory, which is defective in both TgDyrk1a and Dp1Yah models. Why this dosage effect is restricted to recognition memory remains speculative. We may hypothesize that *Cbs* and *Dyrk1a* overdosage only interact in specific regions of the adult brain involved in object discrimination explaining why the increased locomotor activity and the working and visuo-spatial phenotypes induced in Tg(*Dyrk1a*) animals are not affected. Alternatively, objects recognition deficit is likely to result from an impact of DYRK1A on adult brain function while the other phenotypes are the result of an impact during earlier stage of brain development. On the one hand, object recognition has been shown to require undamaged hippocampal perforant path connecting ento/perirhinal cortex with the dentate gyrus for long retention intervals (> 15 min) in rat (73-78). On the other hand, synaptic exchanges between the median prefrontal cortex (mPFC) and the hippocampus seems to be sufficient to support the processing of short-term memory such as working memory observed in the Y maze (79, 80) and hyperactivity is associated with the prefrontal cortex, basal ganglia and cerebellum (81-84). Moreover, long-term recognition memory has been shown to appear in the rat at weaning (post-natal day 21 in the mouse), (85), a period corresponding to the end of neurogenesis and synaptogenesis in the dentate of the hippocampus, and reflecting the general observation of ‘infantile amnesia’ observed on long-term memory tasks but not on short-term memory ability (86).

Our proposal go farther than the demonstration by Zhang et al (23) that the Hsa21 homologous region on the Mmu17 is a key determinant cognitive deficits in DS mouse models. We showed here that CBS is a key gene for DS related phenotypes in mice with the other homologous interval *Cbr3-Fam3b* located on Mmu16, encompassing *Dyrk1a*. We should also consider that in people with DS, both genes are trisomic and thus the recognition memory deficit observed in DS persons and in the complete T21 mouse model (87) certainly depends not only on the interplay between DYRK1A and CBS but also on interaction with other Hsa21 genes that may affect different pathways or different parts of the brain.

The molecular mechanisms involved in *Cbs-Dyr*k1a genetic interaction have been investigated through a quantitative proteomic approach. Although limited due to the complexity of the hippocampus, the results highlight proteins networks interactions between the two trisomic regions. 208 proteins were found deregulated, corresponding to 148 GO categories and pathways, with 72 specific to Dp1Yah (out of 121) and 9 to *Dyrk1a* transgenic model (out of 32; Supplementary table 3) and 5 common to both Dp1Yah and Tg(*Dyrk1a*). More interestingly, GO terms such as cortical cytoskeleton or cytoskeletal protein binding were respectively affected in Dp1Yah and in the Tg(Dyrk1a) but were restored in the double transgenic animals, unravelling somehow the nature of the pathways controlled by the epistatic interaction between CBS and DYRK1A overdosage. DYRK1A is found mainly associated to and modulates the actin cytoskeleton (88). CBS is the major enzyme involved in H2S production in the central nervous system (67). Interestingly increase of H2S activates RAC1 leading to rearrangement of actin cytoskeleton during endothelial cell migration (89). Thus a simple hypothesis would be that the overdosage of CBS will lead to increased H2S production and further activation of RAC1 with effect on actin cytoskeleton rearrangement, a key mechanism involved in synaptic transmission. Remarkably DYRK1A interacts with p120-Catenin-Kaiso and can then modulate Rac1 (90). Thus one working hypothesis is based on CBS and DYRK1A pathways connected through RAC1.

DYRK1A is the main driver of defects in DS mouse models for the homologous region to Hsa21 located on Mmu16 (55). Based on study done in DS models for the Mmu16 homologous region (91), DYRK1A has been selected as a drug target. As reported previously, a treatment with epigallocatechin-3-gallate (EGCG), an inhibitor of DYRK1A kinase activity, can restore some cognitive aspects found altered in people with DS but the gain was limited (92, 93). Nevertheless our results, by adding CBS to the limited number of DS therapeutic targets, may improve the efficiency of DS treatment, in particular by combining multiple therapies for improving the life of DS patients. Finally, an important point to emphasize is that, for DYRK1A as well as for CBS, both loss of function mutations and overdosage lead to intellectual deficiencies. This is important to keep in mind when considering pharmacological intervention that aims at inhibiting one or the other, or both, of these enzymes. Therefore, drug treatment that lead to only a mild inhibition of CBS and/or DYRK1A should be favoured.

## ACKNOWLEDGEMENTS

We thank Dr David Patterson for granting access to the 60.4P102D1 transgenic mice, and Dr. Nathalie Janel for providing the CBS KO mice, Dr. Henri Blehaut for his initial support on the study and the Fondation Jerome Lejeune for making the transgenic line available and their support. We are grateful to members of the research group, of the proteomic platform of the IGBMC laboratory, and of the Mouse Clinical institute (MCI-ICS) for their help and helpful discussion during the project. The project was supported by the French National Centre for Scientific Research (CNRS), the French National Institute of Health and Medical Research (INSERM), the ITMO (“Institut Thématique Multiorganisme”) BCDE (“Biologie Cellulaire, Développement & Evolution”), the University of Strasbourg and the “Centre Europeen de Recherche en Biomedecine”, the “Fondation Jerome Lejeune” and the French state funds through the “Agence Nationale de la Recherche” under the frame programme Investissements d’Avenir labelled (ANR-10-IDEX-0002-02, ANR-10-LABX-0030-INRT, ANR-10-INBS-07 PHENOMIN). The funders had no role in study design, data collection and analysis, decision to publish, or preparation of the manuscript.

## REFERENCES

1. Korbel JO, et al. (2009) The genetic architecture of Down syndrome phenotypes revealed by high-resolution analysis of human segmental trisomies. Proceedings of the National Academy of Sciences of the United States of America 106(29):12031–12036.

2. Lyle R, et al. (2009) Genotype-phenotype correlations in Down syndrome identified by array CGH in 30 cases of partial trisomy and partial monosomy chromosome 21. European Journal of Human Genetics 17(4):454–466.

3. Reeves RH, et al. (1995) A MOUSE MODEL FOR DOWN-SYNDROME EXHIBITS LEARNING AND BEHAVIOR DEFICITS. Nature Genetics 11(2):177–184.

4. Yu T, et al. (2010) A mouse model of Down syndrome trisomic for all human chromosome 21 syntenic regions. Human Molecular Genetics 19(14):2780–2791.

5. Duchon A, et al. (2011) The telomeric part of the human chromosome 21 from Cstb to Prmt2 is not necessary for the locomotor and short-term memory deficits observed in the Tc1 mouse model of Down syndrome. Behavioural Brain Research 217(2):271–281.

6. Glahn DC, Thompson PM, & Blangero J (2007) Neuroimaging endophenotypes: strategies for finding genes influencing brain structure and function. Hum Brain Mapp 28(6):488–501.

7. Brault V, et al. (2015) Opposite phenotypes of muscle strength and locomotor function in mouse models of partial trisomy and monosomy 21 for the proximal Hspa13-App region. PLoS Genet 11(3):e1005062.

8. Herault Y, Duchon A, Velot E, Maréchal D, & Brault V (2012) The in vivo Down syndrome genomic library in mouse. Prog Brain Res 197:169–197.

9. Jiang X, et al. (2015) Genetic dissection of the Down syndrome critical region. Hum Mol Genet.

10. Hall JH, et al. (2016) Tc1 mouse model of trisomy-21 dissociates properties of short-and longterm recognition memory. Neurobiol Learn Mem 130:118–128.

11. Lana-Elola E, et al. (2016) Genetic dissection of Down syndrome-associated congenital heart defects using a new mouse mapping panel. Elife 5.

12. Salehi A, et al. (2006) Increased App expression in a mouse model of Down’s syndrome disrupts NGF transport and causes cholinergic neuron degeneration. Neuron 51(1):29–42.

13. García-Cerro S, et al. (2014) Overexpression of Dyrk1A is implicated in several cognitive, electrophysiological and neuromorphological alterations found in a mouse model of Down syndrome. PLoS One 9(9):e106572.

14. Altafaj X, et al. (2013) Normalization of Dyrk1A expression by AAV2/1-shDyrk1A attenuates hippocampal-dependent defects in the Ts65Dn mouse model of Down syndrome. Neurobiol Dis 52:117–127.

15. Guedj F, et al. (2009) Green tea polyphenols rescue of brain defects induced by overexpression of DYRK1A. PLoS One 4(2):e4606.

16. De la Torre R, et al. (2014) Epigallocatechin-3-gallate, a DYRK1A inhibitor, rescues cognitive deficits in Down syndrome mouse models and in humans. Mol Nutr Food Res 58(2):278–288.

17. de la Torre R, et al. (2016) Safety and efficacy of cognitive training plus epigallocatechin-3-gallate in young adults with Down’s syndrome (TESDAD): a double-blind, randomised, placebo-controlled, phase 2 trial. Lancet Neurol 15(8):801–810.

18. Kim H, et al. (2016) A chemical with proven clinical safety rescues Down-syndrome-related phenotypes in through DYRK1A inhibition. Dis Model Mech 9(8):839–848.

19. Nakano-Kobayashi A, et al. (2017) Prenatal neurogenesis induction therapy normalizes brain structure and function in Down syndrome mice. Proc Natl Acad Sci U S A 114(38):10268–10273.

20. Neumann F, et al. (2018) DYRK1A inhibition and cognitive rescue in a Down syndrome mouse model are induced by new fluoro-DANDY derivatives. Sci Rep 8(1):2859.

21. Pereira PL, et al. (2009) A new mouse model for the trisomy of the Abcg1-U2af1 region reveals the complexity of the combinatorial genetic code of down syndrome. Human Molecular Genetics 18(24):4756–4769.

22. Yu T, et al. (2010) Effects of individual segmental trisomies of human chromosome 21 syntenic regions on hippocampal long-term potentiation and cognitive behaviors in mice. Brain Research 1366:162–171.

23. Zhang L, et al. (2014) Human chromosome 21 orthologous region on mouse chromosome 17 is a major determinant of Down syndrome-related developmental cognitive deficits. Hum Mol Genet 23(3):578–589.

24. Sahún I, et al. (2014) Cognition and Hippocampal Plasticity in the Mouse Is Altered by Monosomy of a Genomic Region Implicated in Down Syndrome. Genetics 197(3):899–912.

25. Marechal D, Lopes Pereira P, Duchon A, & Herault Y (2015) Dosage of the Abcg1-U2af1 region modifies locomotor and cognitive deficits observed in the Tc1 mouse model of Down syndrome. PLoS One 10(2):e0115302.

26. Kimura H (2011) Hydrogen sulfide: its production, release and functions. Amino Acids 41(1):113–121.

27. Chen X, Jhee KH, & Kruger WD (2004) Production of the neuromodulator H2S by cystathionine beta-synthase via the condensation of cysteine and homocysteine. J Biol Chem 279(50):52082–52086.

28. Kamat PK, Kalani A, & Tyagi N (2015) Role of hydrogen sulfide in brain synaptic remodeling. Methods Enzymol 555:207–229.

29. Watanabe M, et al. (1995) MICE DEFICIENT IN CYSTATHIONINE BETA-SYNTHASE-ANIMAL-MODELS FOR MILD AND SEVERE HOMOCYST(E)INEMIA. Proceedings of the National Academy of Sciences of the United States of America 92(5):1585–1589.

30. Butler C, Knox AJ, Bowersox J, Forbes S, & Patterson D (2006) The production of transgenic mice expressing human cystathionine beta-synthase to study Down syndrome. Behav Genet 36(3):429–438.

31. Mantamadiotis T, et al. (2002) Disruption of CREB function in brain leads to neurodegeneration. Nat Genet 31(1):47–54.

32. Guedj F, et al. (2012) DYRK1A: a master regulatory protein controlling brain growth. Neurobiol Dis 46(1):190–203.

33. Lopes Pereira P, et al. (2009) A new mouse model for the trisomy of the Abcg1-U2af1 region reveals the complexity of the combinatorial genetic code of down syndrome. Hum Mol Genet 18(24):4756–4769.

34. Mumberg D, Muller R, & Funk M (1995) YEAST VECTORS FOR THE CONTROLLED EXPRESSION OF HETEROLOGOUS PROTEINS IN DIFFERENT GENETIC BACKGROUNDS. Gene 156(1):119–122.

35. Ito H, Fukuda Y, Murata K, & Kimura A (1983) Transformation of intact yeast cells treated with alkali cations. J Bacteriol 153(1):163–168.

36. Kim AK & Souza-Formigoni MLO (2010) Disulfiram impairs the development of behavioural sensitization to the stimulant effect of ethanol. Behavioural Brain Research 207(2):441–446.

37. Karp NA, et al. (2015) Applying the ARRIVE Guidelines to an In Vivo Database. PLoS Biol 13(5):e1002151.

38. Kilkenny C, Browne WJ, Cuthill IC, Emerson M, & Altman DG (2010) Improving bioscience research reporting: the ARRIVE guidelines for reporting animal research. PLoS Biol 8(6):e1000412.

39. Asimakopoulou A, et al. (2013) Selectivity of commonly used pharmacological inhibitors for cystathionine beta synthase (CBS) and cystathionine gamma lyase (CSE). Br J Pharmacol 169(4):922–932.

40. Thorson MK, et al. (2015) Marine natural products as inhibitors of cystathionine beta-synthase activity. Bioorg Med Chem Lett 25(5):1064–1066.

41. Thorson MK, Majtan T, Kraus JP, & Barrios AM (2013) Identification of cystathionine β-synthase inhibitors using a hydrogen sulfide selective probe. Angew Chem Int Ed Engl 52(17):4641–4644.

42. Zhou Y, et al. (2013) High-throughput tandem-microwell assay identifies inhibitors of the hydrogen sulfide signaling pathway. Chem Commun (Camb) 49(100):11782–11784.

43. Chao C, et al. (2016) Cystathionine-beta-synthase inhibition for colon cancer: Enhancement of the efficacy of aminooxyacetic acid via the prodrug approach. Mol Med 22.

44. Druzhyna N, et al. (2016) Screening of a composite library of clinically used drugs and well-characterized pharmacological compounds for cystathionine β-synthase inhibition identifies benserazide as a drug potentially suitable for repurposing for the experimental therapy of colon cancer. Pharmacol Res 113(Pt A):18–37.

45. Lasserre JP, et al. (2015) Yeast as a system for modeling mitochondrial disease mechanisms and discovering therapies. Dis Model Mech 8(6):509–526.

46. Voisset C, et al. (2014) A yeast-based assay identifies drugs that interfere with Epstein-Barr virus immune evasion. Dis Model Mech.

47. Couplan E, et al. (2011) A yeast-based assay identifies drugs active against human mitochondrial disorders. Proc Natl Acad Sci U S A 108(29):11989–11994.

48. Khurana V, Tardiff DF, Chung CY, & Lindquist S (2015) Toward stem cell-based phenotypic screens for neurodegenerative diseases. Nat Rev Neurol 11(6):339–350.

49. Bach S, et al. (2003) Isolation of drugs active against mammalian prions using a yeast-based screening assay. Nat Biotechnol 21(9):1075–1081.

50. Khurana V & Lindquist S (2010) OPINION Modelling neurodegeneration in Saccharomyces cerevisiae: why cook with baker’s yeast? Nature Reviews Neuroscience 11(6):436–449.

51. Lista MJ, et al. (2017) Nucleolin directly mediates Epstein-Barr virus immune evasion through binding to G-quadruplexes of EBNA1 mRNA. Nature Communications 8.

52. Mayfield JA, et al. (2012) Surrogate genetics and metabolic profiling for characterization of human disease alleles. Genetics 190(4):1309–1323.

53. Kruger WD & Cox DR (1994) A yeast system for expression of human cystathionine beta-synthase: structural and functional conservation of the human and yeast genes. Proc Natl Acad Sci U S A 91(14):6614–6618.

54. Aiyar RS, et al. (2014) Mitochondrial protein sorting as a therapeutic target for ATP synthase disorders. Nat Commun 5:5585.

55. Duchon A & Herault Y (2016) DYRK1A, a Dosage-Sensitive Gene Involved in Neurodevelopmental Disorders, Is a Target for Drug Development in Down Syndrome. Front Behav Neurosci 10:104.

56. Hamelet J, et al. (2009) Effect of hyperhomocysteinemia on the protein kinase DYRK1A in liver of mice. Biochem Biophys Res Commun 378(3):673–677.

57. Planque C, et al. (2013) Mice deficient in cystathionine beta synthase display increased Dyrk1A and SAHH activities in brain. J Mol Neurosci 50(1):1–6.

58. Noll C, et al. (2009) DYRK1A, a novel determinant of the methionine-homocysteine cycle in different mouse models overexpressing this Down-syndrome-associated kinase. PLoS One 4(10):e7540.

59. Kwiatkowski TJ, Jr., et al. (2009) Mutations in the FUS/TLS Gene on Chromosome 16 Cause Familial Amyotrophic Lateral Sclerosis. Science 323(5918):1205–1208.

60. Vance C, et al. (2009) Mutations in FUS, an RNA Processing Protein, Cause Familial Amyotrophic Lateral Sclerosis Type 6. Science 323(5918):1208–1211.

61. Polymeropoulos MH, et al. (1997) Mutation in the alpha-synuclein gene identified in families with Parkinson’s disease. Science 276(5321):2045–2047.

62. Spillantini MG, et al. (1997) alpha-synuclein in Lewy bodies. Nature 388(6645):839–840.

63. Sorensen JB, et al. (2003) Differential control of the releasable vesicle pools by SNAP-25 splice variants and SNAP-23. Cell 114(1):75–86.

64. McMahon HT & Sudhof TC (1995) SYNAPTIC CORE COMPLEX OF SYNAPTOBREVIN, SYNTAXIN, AND SNAP25 FORMS HIGH-AFFINITY ALPHA-SNAP FINDING SITE. Journal of Biological Chemistry 270(5):2213–2217.

65. Zhou Q, et al. (2017) The primed SNARE-complexin-synaptotagmin complex for neuronal exocytosis. Nature 548(7668):420–425.

66. Régnier V, et al. (2012) Brain phenotype of transgenic mice overexpressing cystathionine β-synthase. PLoS One 7(1):e29056.

67. Kimura H (2002) Hydrogen sulfide as a neuromodulator. Mol Neurobiol 26(1):13–19.

68. Kimura H (2000) Hydrogen sulfide induces cyclic AMP and modulates the NMDA receptor. Biochem Biophys Res Commun 267(1):129–133.

69. Hu LF, Lu M, Hon Wong PT, & Bian JS (2011) Hydrogen sulfide: neurophysiology and neuropathology. Antioxid Redox Signal 15(2):405–419.

70. Johansson B (1992) A review of the pharmacokinetics and pharmacodynamics of disulfiram and its metabolites. Acta Psychiatr Scand Suppl 369:15–26.

71. Barth KS & Malcolm RJ (2010) Disulfiram: an old therapeutic with new applications. CNS Neurol Disord Drug Targets 9(1):5–12.

72. Delabar JM, et al. (2014) One-carbon cycle alterations induced by Dyrk1a dosage. Mol Genet Metab Rep 1:487–492.

73. Antunes M & Biala G (2012) The novel object recognition memory: neurobiology, test procedure, and its modifications. Cognitive Processing 13(2):93–110.

74. Clark RE, Zola SM, & Squire LR (2000) Impaired recognition memory in rats after damage to the hippocampus. Journal of Neuroscience 20(23):8853–8860.

75. Clarke JR, Cammarota M, Gruart A, Izquierdo I, & Delgado-Garcia JM (2010) Plastic modifications induced by object recognition memory processing. Proceedings of the National Academy of Sciences of the United States of America 107(6):2652–2657.

76. Reger ML, Hovda DA, & Giza CC (2009) Ontogeny of Rat Recognition Memory Measured by the Novel Object Recognition Task. Developmental Psychobiology 51(8):672–678.

77. Stackman RW, Cohen SJ, Lora JC, & Rios LM (2016) Temporary inactivation reveals that the CA1 region of the mouse dorsal hippocampus plays an equivalent role in the retrieval of longterm object memory and spatial memory. Neurobiology of Learning and Memory 133:118–128.

78. Warburton EC & Brown MW (2015) Neural circuitry for rat recognition memory. Behavioural Brain Research 285:131–139.

79. Benchenane K, et al. (2010) Coherent theta oscillations and reorganization of spike timing in the hippocampal-prefrontal network upon learning. Neuron 66(6):921–936.

80. Wei J, Bai WW, Liu TT, & Tian X (2015) Functional connectivity changes during a working memory task in rat via NMF analysis. Frontiers in Behavioral Neuroscience 9.

81. Lin JD, et al. (2004) Defects in adaptive energy metabolism with CNS-Linked hyperactivity in PGC-1 alpha null mice. Cell 119(1):121-+.

82. Vallone D, Picetti R, & Borrelli E (2000) Structure and function of dopamine receptors. Neuroscience and Biobehavioral Reviews 24(1):125–132.

83. Bymaster FP, et al. (2002) Atomoxetine increases extracellular levels of norepinephrine and dopamine in prefrontal cortex of rat: A potential mechanism for efficacy in Attention Deficit/Hyperactivity Disorder. Neuropsychopharmacology 27(5):699–711.

84. Cador M, Robbins TW, & Everitt BJ (1989) INVOLVEMENT OF THE AMYGDALA IN STIMULUS REWARD ASSOCIATIONS-INTERACTION WITH THE VENTRAL STRIATUM. Neuroscience 30(1):77–86.

85. Anderson MJ, et al. (2004) Effects of ontogeny on performance of rats in a novel object-recognition task. Psychological Reports 94(2):437–443.

86. Rudy JW & Morledge P (1994) ONTOGENY OF CONTEXTUAL FEAR CONDITIONING IN RATS-IMPLICATIONS FOR CONSOLIDATION, INFANTILE AMNESIA, AND HIPPOCAMPAL SYSTEM FUNCTION. Behavioral Neuroscience 108(2):227–234.

87. Belichenko PV, et al. (2015) Down Syndrome Cognitive Phenotypes Modeled in Mice Trisomic for All HSA 21 Homologues. Plos One 10(7).

88. Park J, Sung JY, Song WJ, Chang S, & Chung KC (2012) Dyrk1A negatively regulates the actin cytoskeleton through threonine phosphorylation of N-WASP. J Cell Sci 125(Pt 1):67–80.

89. Zhang LJ, Tao BB, Wang MJ, Jin HM, & Zhu YC (2012) PI3K p110α isoform-dependent Rho GTPase Rac1 activation mediates H2S-promoted endothelial cell migration via actin cytoskeleton reorganization. PLoS One 7(9):e44590.

90. Hong JY, et al. (2012) Down’s-syndrome-related kinase Dyrk1A modulates the p120-catenin-Kaiso trajectory of the Wnt signaling pathway. J Cell Sci 125(Pt 3):561–569.

91. Herault Y, et al. (2017) Rodent models in Down syndrome research: impact and future opportunities. Dis Model Mech 10(10):1165–1186.

92. de la Torre R, et al. (2016) Safety and efficacy of cognitive training plus epigallocatechin-3-gallate in young adults with Down’s syndrome (TESDAD): a double-blind, randomised, placebo-controlled, phase 2 trial. Lancet Neurology 15(8):801–810.

93. De la Torre R, et al. (2014) Epigallocatechin-3-gallate, a DYRK1A inhibitor, rescues cognitive deficits in Down syndrome mouse models and in humans. Molecular Nutrition & Food Research 58(2):278–288.

